# Linear morphometrics fail to support strong sexual dimorphism in *Uintatherium anceps*

**DOI:** 10.64898/2026.03.05.709752

**Authors:** Kevin D. Mulcahy

## Abstract

Uintatheres, mammals belonging to the extinct order Dinocerata, are among the most recognizable of all Paleogene (∼66 – 23 Ma) organisms. Unmistakable for their bizarre skulls with multiple pairs of horns and saber-like upper canines, uintatheres have captivated paleontologists since the late nineteenth century. Since their initial discovery, uintatheres have been regarded as a classic example of dramatic sexual dimorphism in the fossil record, with males purported to be larger and possess more prominent horns and canines than females. However, the hypothesis that uintatheres were highly sexually dimorphic has never been formally tested. Here, I use traditional, linear morphometrics on a collection including most known skulls of *Uintatherium anceps* to quantify patterns of cranial variation within this taxon. Despite using a variety of traditional and novel statistical methods, I fail to detect any evidence of strong sexual dimorphism in *Uintatherium*. To verify my approach, I assembled a similarly sized dataset from *Bison bison* as an extant analog, and found strong, consistent evidence of sexual dimorphism. In light of these findings, as well as the current understanding of uintathere systematics and paleoecology, I argue that strong sexual dimorphism should not be treated as the null hypothesis for this clade.

## Introduction

Sexual dimorphism is the phenomenon in which individuals of different sexes of the same species differ in their morphology beyond primary sexual characteristics [1–5]. Naturalists have observed patterns of sexual dimorphism for centuries, and description of dimorphism and its causes factored heavily into much of Darwin’s work on sexual selection [1,2]. Among mammals, researchers historically accepted that males tended to be larger and possess more elaborate secondary sexual characteristics than females [1,2,6–10]. This notion was explained by higher investment in reproduction by females, and therefore higher selective pressures and competition among males, especially in polygynous mating systems [2,6,10–13]. Indeed, in many of the most famous and well-studied extant mammals, males are significantly larger than females and possess obvious secondary sexual characters, allowing for near-intuitive sex identification (e.g., [2,14–16]).

However, while sexual dimorphism has been conclusively documented in many taxa, advances in mammalian research have demonstrated that patterns of dimorphism are not ubiquitous or uniform among mammals. In the 1970s, Ralls reviewed rates of sexual size dimorphism across mammalian clades and found that species with little dimorphism or larger females were surprisingly numerous [17,18]. Building on Ralls work, in 2024, Tombak and colleagues surveyed extant placental mammals and found that males are not in fact measurably larger than females in most species [10]. Furthermore, rates of sexual size dimorphism have been shown to carry strong phylogenetic signal, with ordinal level affinities often predicting patterns of dimorphism [10,19]. For instance, while species of artiodactyls and primates almost universally exhibit larger males than females, monomorphism and larger females are common among rodents and bats [10,11,20–23]. It has therefore become increasingly unclear whether classical larger-male patterns of sexual dimorphism were ancestral among most mammals.

Among fossil vertebrates, sexual dimorphism has a rich history of documentation and research. Especially in the field of paleoanthropology, the study of purported patterns of sexual dimorphism in the fossil record has formed a key component of research into the evolution of sexual selection and social behavior more broadly [20,24–26]. Instances of sexual dimorphism have been reported among extinct members of virtually all major vertebrate clades, including mammals, birds, and nonavian dinosaurs (e.g., [27–39]). However, researchers have also raised caution in attributing patterns of morphological variation among extinct taxa to sexual dimorphism [40–44]. When working with extant animals, it is relatively easy to test for and quantify patterns of sexual dimorphism, as males and females can usually be observationally or experimentally distinguished and therefore compared [45,46]. In contrast, even in truly dimorphic taxa, fossilized remains rarely preserve the primary sexual characters necessary to definitively determine the sex of an individual [47,48]. This lack of a priori knowledge of individual sex precludes the use of classic statistical tests and quantitative analyses of dimorphism, as patterns of morphological variation could instead represent ontogeny, taxonomic differences, phenotypic plasticity, or random individual variation [40,43,49–51]. Additional factors such as small sample sizes and potentially biased sex ratios present further challenges for the study of sexual dimorphism in the fossil record [40,43,48,52].

Still, approaches to detect and quantify patterns of sexual dimorphism in the fossil record have been developed and applied. One relatively intuitive approach to detect patterns of dimorphism is to test the distribution of a feature of interest for statistically significant deviation from normality or unimodality [43,53,54]. However, this approach has been criticized for high type I and type II error rates, depending on the size and true sex-ratio of the sample. Other methods, including mixture modeling, Principal Component Analysis (PCA), and cluster analysis have been advocated as potentially more robust approaches to detect underlying patterns of dimorphism among traits of interest in the fossil record [43,54–58]. Still other novel frameworks to detect sexual dimorphism in the fossil record continue to be developed [59].

One group of fossil animals for which sexual dimorphism has been widely accepted are the uintatheres, extinct mammals belonging to the order Dinocerata [60–65]. Uintatheres ranged from the late Paleocene to the middle Eocene (∼60-40 Ma) in Asia and North America, and were graviportal, vaguely ungulate-like eutherians, representing the first truly massive land mammals, with body masses potentially exceeding 4,500 kg [60,64,66,67]. Uintatheres, especially the derived uintatheriines, are easily recognized by their bizarre skulls, with later forms possessing three pairs of cranial protuberances and massive, saber-like canines (Fig 1) [60,62,65]. Likely due to their massive body size and enigmatic appearance, uintatheres have figured heavily in the history of American paleontology, serving as key taxa in the infamous “Bone Wars” of Marsh and Cope [60,68,69]. Despite such a colorful history of research, the broader affinities of uintatheres remain unresolved, with various hypotheses aligning them with Laurasiatheria, Euarchontoglires, Afrotheria, or other extinct and early diverging mammalian clades such as Pantodonta [60,63,68,70].

**Figure 1.**
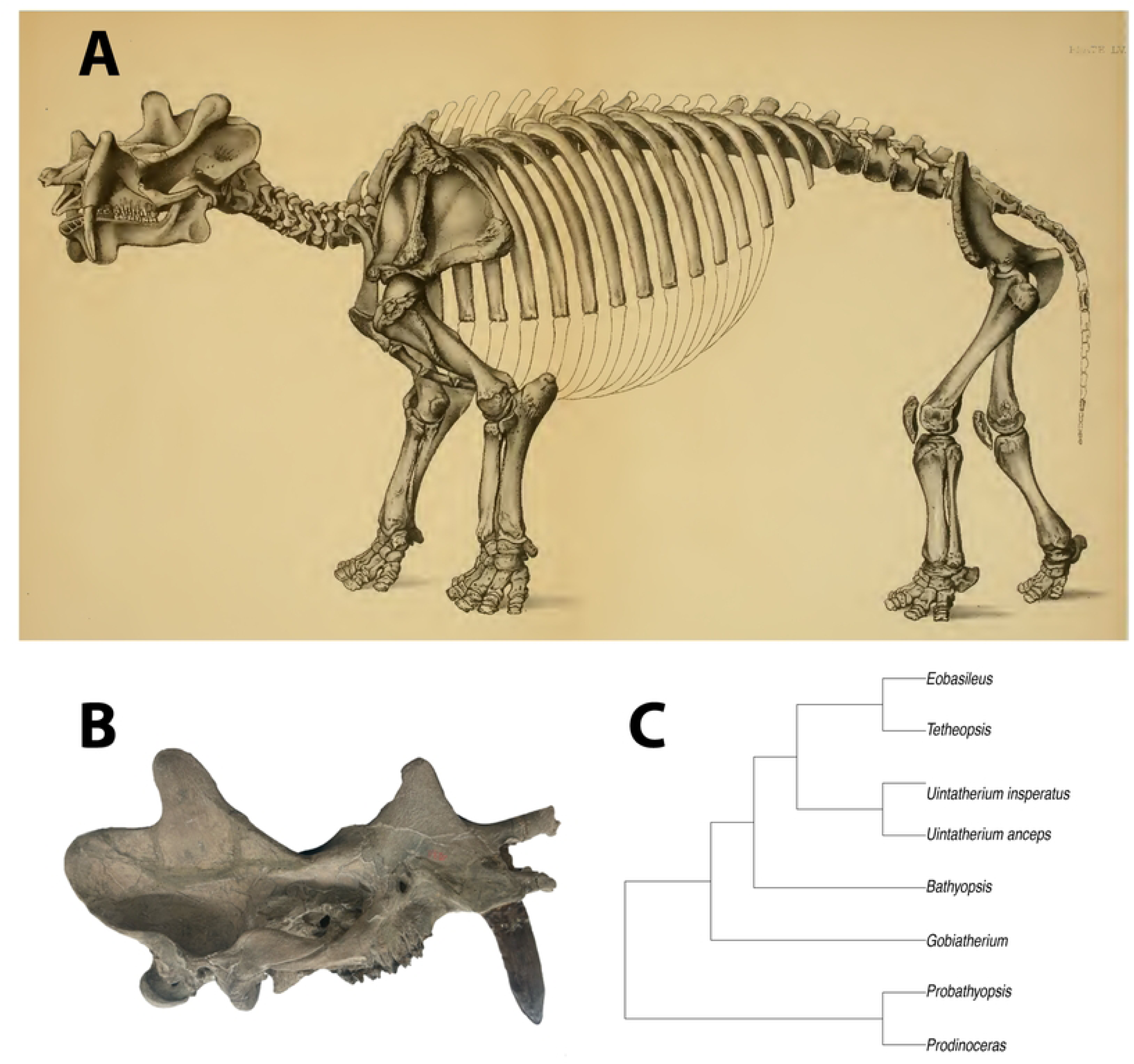
Osteology and systematics of Dinocerata. (**A**) Plate LV from Marsh [60] showing a restoration of *Uintatherium anceps* (Marsh’s holotype of *Dinoceras laticeps*, YPM 11036). (**B**) Skull of *Uintatherium anceps* (YPM 11036). (**C**) Proposed phylogenetic relationships within Dinocerata, adapted from Schoch and Lucas [63]. **Abbreviations**: YPM, Yale Peabody Museum of Natural History.

In 1886, Marsh first posited that uintatheres were highly dimorphic, and this is one of the earliest reports of sexual dimorphism in the entire history of vertebrate paleontology [60]. Subsequent authors have expanded on Marsh’s initial claims and attributed the wide range of cranial variation observed within uintathere species to significant differences between sexes (Fig 2) [62,63,65]. Traditionally, large uintathere specimens with prominent horns and canines have been interpreted as male, while smaller individuals with less dramatic craniodental features have been identified as females [60,62,63,65]. However, to date, the hypothesis that the skulls of uintatheres display a high degree of sexual dimorphism has not been adequately tested or quantitatively validated. Furthermore, while previous authors have agreed that uintatheres were strongly dimorphic, they have differed in their interpretation as to how the sexes actually varied. For example, one particularly small and gracile skull of *Uintatherium anceps*, has been interpreted as both a deformed female and an immature male [62,65]. Additionally, authors have also differed in their interpretations of which uintathere cranial features were dimorphic. For instance, Marsh claims that females of *U. anceps* lacked the prominent inframandibular flange seen in males, while Wheeler argues for the presence of fully developed flanges in individuals of both sexes [60,62].

**Figure 2.**
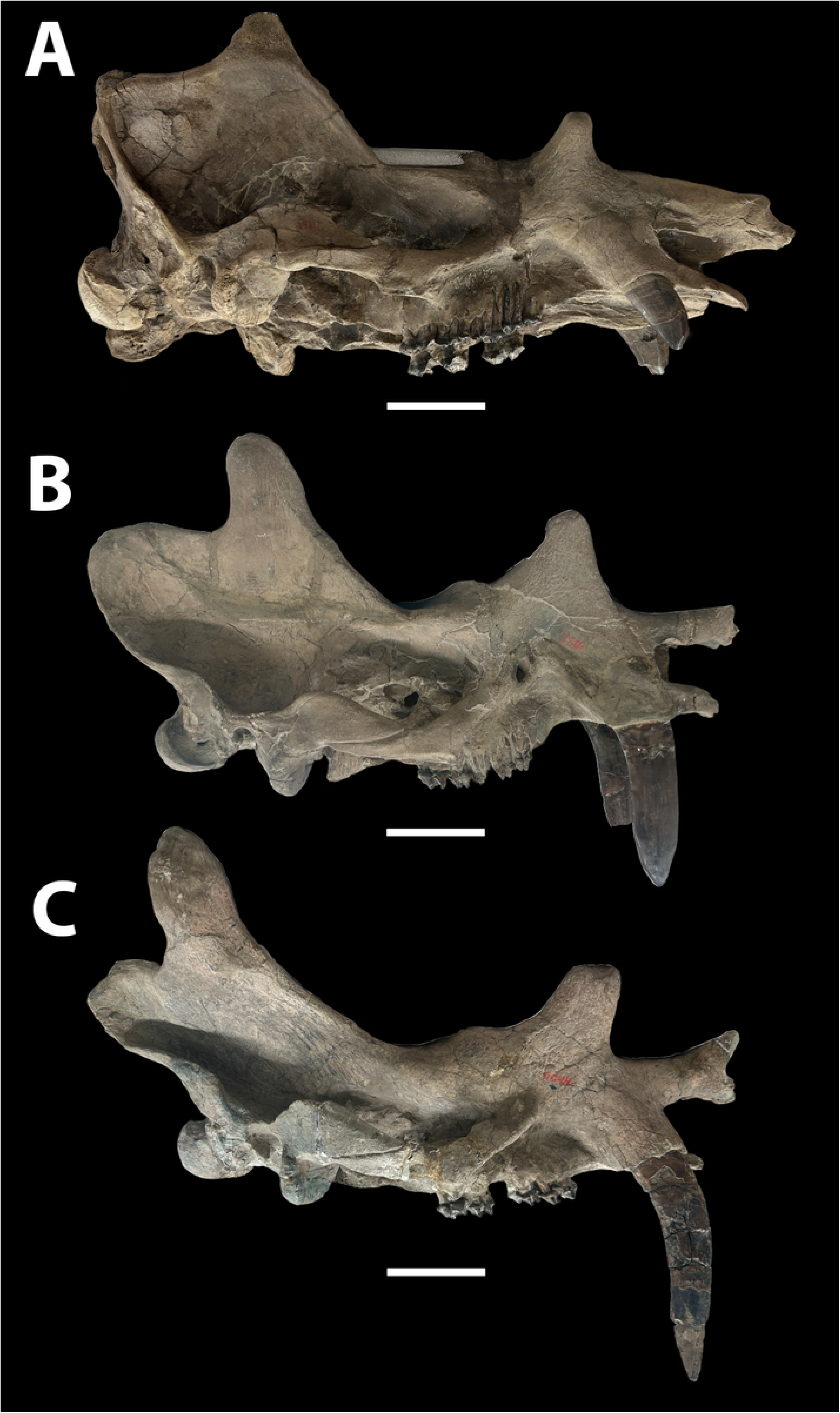
Cranial variation in the skulls of *Uintatherium anceps*. (**A**) AMNH 1671, purported female according to Wheeler [62] and Turnbull [65]. (**B**) YPM 11036, sex unreported. (**C**) YPM 11044, purported male according to Marsh [60]. Scalebars equal 10 cm. **Abbreviations**: AMNH, American Museum of Natural History; YPM, Yale Peabody Museum of Natural History.

Given their early divergence and potentially basal phylogenetic position, determining whether uintatheres were sexually dimorphic could yield insights into the character polarity of sexual dimorphism among broader mammalian clades. Here, I present a quantitative morphometric analysis of cranial morphology in uintatheres and perform the first tests of the hypothesis that they were sexually dimorphic. As previous authors have expressed skepticism towards traditional approaches used to detect patterns of sexual dimorphism in fossil taxa, I employ a variety of methods including classical tests of univariate normality and unimodality, principal component analysis, cluster analysis, and a novel approach based on effect size statistics [40,43,50,59]. To validate my approach, I gathered a novel dataset from skulls of the American bison as an extant dimorphic analog and performed the same analyses. I believe the following represents a robust analysis of purported patterns of sexual dimorphism in the fossil record which can serve as a reference for students of other taxa.

## Materials and Methods

### Collections and permissions

Specimens referenced in this study are curated in the following collections: **AMNH**, American Museum of Natural History, New York, New York, USA; **YPM**, Yale Peabody Museum of Natural History, New Haven, Connecticut, USA; **YPM-PU**, Princeton University collections acquired by the Yale Peabody Museum, New Haven, Connecticut, USA; **PM**, Princeton Museum specimens now housed at the Field Museum of Natural History, Chicago, Illinois, USA; **UW**, University of Wyoming Geological Museum, Laramie, Wyoming, USA; **KU:Mamm**, mammalogy collections at the University of Kansas Natural History Museum, Lawrence, Kansas, USA. All necessary permissions were obtained for the described study, which complied with all relevant regulations.

### Specimens and cranial measurements

For this study, I analyzed 27 complete and partial skulls of mature individuals (possessing full adult dentition) of *Uintatherium anceps* at the American Museum of Natural History and the Yale Peabody Museum of Natural History. *U. anceps* was chosen as the focal taxon as it is by far the best represented uintatheriine. In addition, data from two mature skulls at the Field Museum of Natural History were integrated from Turnbull [65]. Together, this dataset encompasses nearly all *U. anceps* crania known. When assessing evidence of sexual dimorphism, certain cranial features were of particular interest, including the anteriorly projecting premaxillaries and nasals (and prenasal bones on some specimens), the three pairs of horns or cranial protuberances (nasal, maxillary, and parietal), the posteriorly raised occipital crest, and the saber-like canines (Fig 3A).

**Figure 3.**
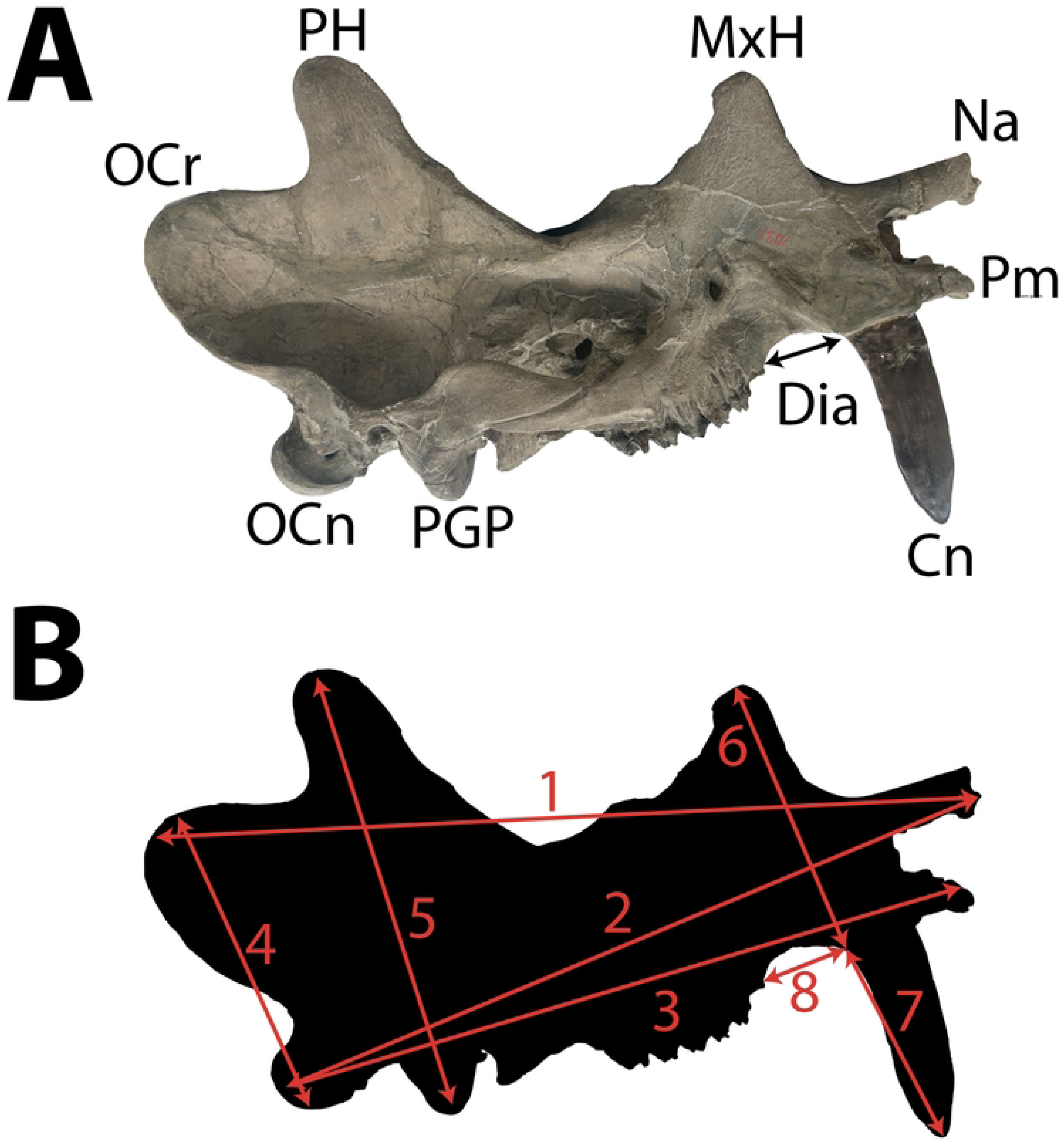
Cranial features and measurements referenced in study of *Uintatherium anceps*. (**A**) Relevant anatomical features on the skull of *Uintatherium anceps* (YPM 11036): OCr, Occipital crest; OCn, Occipital condyle; PGP, Postglenoid process; PH, Parietal horn/protuberance; MxH, Maxillary horn/protuberance; Na, Anterior extent of nasals; Pm, anterior extent of premaxillaries; Cn, Canine; Dia, Diastema. (**B**) Linear cranial metrics used in analyses: 1, Length of skull from nasals to occipital crest; 2, Length of skull from nasals to occipital condyles; 3, Length of skull from premaxillaries to occipital condyles; 4, Height of skull from occipital condyle to occipital crest; 5, Height of parietal protuberance; 6, Height of maxillary protuberance; 7, Length of canine; 8, Diastemal length. **Abbreviations**: YPM, Yale Peabody Museum of Natural History.

The large size of these skulls has precluded computed tomography (CT) scanning and therefore three-dimensional geometric morphometric analyses. Furthermore, due to their tremendous mass, age (both geological and historical), and fragile reconstruction with plaster, I wanted to physically manipulate the skulls as little as possible. Specimens at the AMNH and YPM are firmly affixed to plaster or wooden bases, which allow the fossils to be easily moved between shelves but largely prohibit any rotation or free motion of the skull itself. As some skulls are oriented with the dorsal side facing upwards and others are oriented with the ventral side up, I chose to employ a traditional morphometric approach, through which patterns of morphological variation were captured by linear measurements of the skull that could be observed in lateral view. Therefore, data used in analyses was gathered by taking images of the skulls in lateral view and measuring the distance between landmarks of interest in the software ImageJ.

Eight cranial metrics were chosen to be measured for each skull, capturing both the overall size of each specimen as well as the relative development of the purportedly dimorphic characters (Fig 3B and S1 Table). These metrics were adapted from Turnbull, allowing for direct integration of the FMNH specimens. While this linear morphometric approach likely limits statistical power in comparison to two- or three-dimensional geometric morphometrics, it allows for inclusion of partial and damaged skulls in the analysis. Furthermore, by incorporating measurements drawn on by Turnbull in his discussion of sexual dimorphism in uintatheres, this approach allows me to more directly test previous interpretations of the role of sexual dimorphism in shaping morphological variation in the clade. Linear morphometrics can also be more easily applied in analyses of purported patterns of dimorphism in other clades with similarly problematic fossil records, thus promoting broader reproducibility and comparison.

Measurements obtained in ImageJ were preferred to those gathered by hand, as the large size, varying degrees of preservation, and inconsistent storage conditions precluded consistency in manual measurements across specimens. Some degree of lens distortion likely affects the accuracy of linear measurements obtained from two-dimensional images, but as the same methodology was used for all specimens, error should be relatively consistent across individuals and largely inconsequential to the results of subsequent analyses. To verify that any error in ImageJ measurements was systematic, Bland-Altman plots were generated and tests for association using Pearson’s product moment correlation coefficient were performed to confirm that the difference between the by-hand and ImageJ values did not covary with the size of the skull (S1 Fig and S2 Table). Results were non-significant in all eight tests, indicating that the margin of measurement error did not increase with specimen size.

### Traditional approaches to detect sexual dimorphism

To assess evidence of sexual dimorphism, I employed a variety of statistical approaches, all using R statistical software. First, simple histograms and violin plots were generated to visualize the distribution of each cranial metric, using the plotting tools available in base R and ‘ggplot2’ [71,72]. I then performed Shapiro-Wilk tests of normality and Hartigan’s dip tests of unimodality on the distributions of each metric, using base R and the package ‘diptest’ [73]. Dip test p-values were computed through Monte Carlo simulation with 2,000 replicates. To further assess evidence of multimodality, I also performed univariate mixture modeling, using the package ‘mclust’ to determine how many underlying normal distributions (Gaussians) best fit the data structure for each cranial metric [74]. For each metric, the model with the optimal number of clusters was identified based on the Bayesian Information Criterion (BIC) [75]. In principle, if *U. anceps* were sexually dimorphic, I would expect any metric exhibiting dimorphism to be best fit by a model with two components, one for males and one for females. While mixture modeling may still be prone to overfitting in small samples, it has been proposed as a more powerful means of detecting sexual dimorphism in the fossil record than simple tests of normality or unimodality [43,59].

While statistically significant deviation from a normal, unimodal distribution in any one metric may provide evidence of sexual dimorphism, non-significant test results do not preclude it, nor do significant results confirm its presence. Therefore, I also employed several multivariate approaches. First, I performed hierarchical clustering, by which skulls are clustered and arranged in a dendrogram based on similarities and differences across cranial metrics, using the R package ‘ComplexHeatMap’ [76,77]. One advantage of such an approach is that it allows for intuitive assessment of previous authors’ hypotheses as to which specimens represent males as opposed to females. In hierarchical clustering, specimens that share similar scores across cranial metrics should be placed near one another in the resulting dendrogram. As such, if males and females differ consistently in their morphology, one would expect the purported members of each sex to cluster together on the dendrogram. As hierarchical clustering cannot be performed with missing values, imputation was necessary for skulls that were partially broken. Where appropriate (three or fewer values missing for a specimen), imputation was performed with a Principal Components Analysis model using the R package ‘missMDA’ [78]. Particularly fragmentary specimens (more than three missing values) were not considered in this and subsequent multivariate analyses. Before clustering, all values were standardized so that each cranial metric was normalized and weighted equally in multivariate analysis.

Principal component analysis (PCA) is another method that can be used to visualize the distribution of specimens within a multidimensional morphospace. PCA is a common ordination technique by which multiple dimensions of variation are reduced into a small number of orthogonal principal components capturing covariation across multiple metrics [79]. These principal components can then be analyzed for their eigenvalues (how much of the overall variation in the dataset is captured by each principal component) and loadings (how much each variable contributes to each principal component). Similar in principle to hierarchical clustering, if uintatheres differ consistently in their morphology based on sex, one would expect males and females to occupy distinct morphospaces in PCA. PCA was performed using the R package ‘FactoMineR’ and results were visualized in a bivariate plot displaying scores for PC1 and PC2 [80].

Another multivariate method which can be used to explore purported patterns of sexual dimorphism is k-means cluster analysis. Unlike hierarchical clustering, which groups the data into a dendrogram, k-means clustering is partitional, meaning the user specifies how many discrete clusters the data should form [81]. If, as would be expected in a highly dimorphic taxon, male and female uintatheres displayed consistent differences in their overall morphology, splitting the dataset into two clusters might indicate which specimens were males and which were females. I therefore performed k-means clustering to partition the data into two clusters using the ‘stats’ package and superimposed results onto the bivariate plot generated through PCA. This allowed me to compare the morphospaces of the two most favorable clusters with those occupied by purported males and females. If uintatheres exhibited strong sexual dimorphism, I would expect purported males to fall within one cluster while purported females group into the other. Fisher’s exact test was performed to determine if placement of individual skulls into the two most favorable clusters correlated with sex identity as posited by previous authors.

While k-means clustering allows one to partition the data into the two most optimal clusters, there are also multivariate methods that allow for the determination of how many discrete partitions best model the data. Use of these approaches therefore allows one not only to see which skulls tend to cluster together, but also to ask whether the data is well-fit by a bipartition model in the first place. For this study, total within sum of squares, average silhouette width, and gap statistics were used to determine the optimal number of clusters formed by the data using the ‘factoextra’ package [82]. As with univariate mixture modeling, if uintatheres were highly dimorphic, I would expect this dataset to be best fit by a model with two clusters.

### Effect size statistics

I also employed a new approach developed by Saitta et al. [59] to investigate purported patterns of sexual dimorphism in *U. anceps* using effect size statistics. In this framework, the values for a trait of interest are plotted along a growth curve, and residuals from that curve are calculated for each specimen. All specimens with positive residuals are assigned to one sex and all specimens with negative residuals are assigned to the other. The effect size is then calculated as the proportional difference between the asymptotic trait values of the two sexes.

Saitta’s approach is useful in providing a framework in which the magnitude of sexual dimorphism can be estimated without needing to explicitly accept or reject its presence. This design is particularly powerful when considering traits across multiple taxa or multiple traits within the same taxon. For instance, body size should show a much higher effect size in a highly dimorphic taxon than a monomorphic one. Similarly, within a single dimorphic taxon, a secondary sexual characteristic like horn size should show a higher effect size than a relatively monomorphic character. Although Saitta et al. do not argue that residual-based assignments should be used to determine which fossil specimens were male or female, they note that analyses using extant crocodilian and avian datasets were more accurate at predicting sex than would be expected at random [42,59,83,84]. To my knowledge, this study represents the first application of this approach to the study of sexual dimorphism in extinct mammals.

To generate growth curves, a proxy for age in uintatheres was necessary. For adult specimens, I used tooth wear, placing specimens into three grades reflecting relative wear, as has been done in extant herbivores (S3 Table) [85–87]. Based on patterns observed in extant ungulates and estimated lifespans of extinct mammalian herbivores, each grade was assigned hypothetical upper and lower bounds, and the age of each individual specimen was drawn from a uniform distribution within these bounds [85,88,89]. Application of Saitta et al.’s effect based approach requires sampling across the ontogenetic range of the taxon of interest. As the AMNH and YPM only possess skulls of *Uintatherium* from adult or near adult animals, metrics for two juvenile specimens at the FMNH were obtained from Turnbull [65]. These skulls were placed into a separate grade below any of the adult specimens. To account for differences in measurement techniques and lens distortion, Turnbull’s measurements were adjusted based on the observed differences between the by-hand and ImageJ measurements. For each metric, if there was a statistically significant correlation between the by-hand and ImageJ values, Turnbull’s value was incorporated directly into the linear regression of the two, otherwise the average difference between the ImageJ and by-hand values for a metric was added to Turnbull’s value.

I also simulated five newborn specimens to further anchor the growth curves for each metric. To do this, I estimated cranial metrics for PM 55406A, a partial newborn skull reported by Turnbull, by scaling available material to the next smallest juvenile remains. I then simulated each newborn specimen by drawing from uniform distributions between 75% the estimate for PM 55406A and 95% the value for the smallest juvenile specimen. As with the adults, the ages of juvenile and newborn specimens were estimated by drawing from uniform distributions between the hypothetical bounds of their respective age grades (S3 Table).

Gompertz regression curves, commonly used in the study of growth among mammals, were fit to the observed values of each metric to simulate growth curves using the R package, ‘minpack.lm’ [90–92]. Residuals were plotted using the ‘stats’ package, and Gompertz regression curves were then fit to the subsets of specimens with positive and negative residuals. The effect size was quantified as the difference between the horizontal asymptotes of these two groups. Ninety-five percent confidence and prediction intervals for each sex were obtained using the R package, ‘propagate’ [93].

### Extant analogy

To contextualize results and validate my approach, I also performed all analyses in the American bison (*Bison bison*). *B. bison* was chosen as an extant analog because of its large size, herbivorous feeding adaptations, and overall pattern of sexual dimorphism including both body size and the prominence of cranial horns [94–96]. Furthermore, bison skulls retain a bony horn core or cornual process, providing an osteological correlate analogous to the horns of uintatheres [85,94]. Additionally, as *B. bison* was historically abundant across the central United States, the species is represented by extensive museum collections, with enough skulls to facilitate analyses of dimorphism. For this study, I gathered cranial metrics from 23 skulls of *B. bison* at KU:Mamm. Of these, nine were female, seven were male, and seven lacked associated sex metadata.

To maintain consistency with the uintatheres, I chose to measure eight cranial metrics in *B. bison*, capturing both the overall size of the skull and the relative prominence of the horns (Fig 4 and S4 Table). As *Bison* only possess one pair of horns that project laterally, I chose measurements that could all be observed in anterior view. Otherwise, all analyses were performed following the same protocols employed with the uintatheres. For effect size analyses, age estimates were drawn from uniform distributions within Fuller’s age grades based on dental eruption and wear (S5 Table) [85]. Similar to the uintatheres, the bison dataset consisted only of adults, near-adults, and one juvenile. As such, I again simulated five newborn specimens by estimating values based on the available literature [96]. The values of each newborn’s cranial metrics were drawn from uniform distributions set between 75% and 125% of the estimate for each trait derived from the literature.

**Figure 4.**
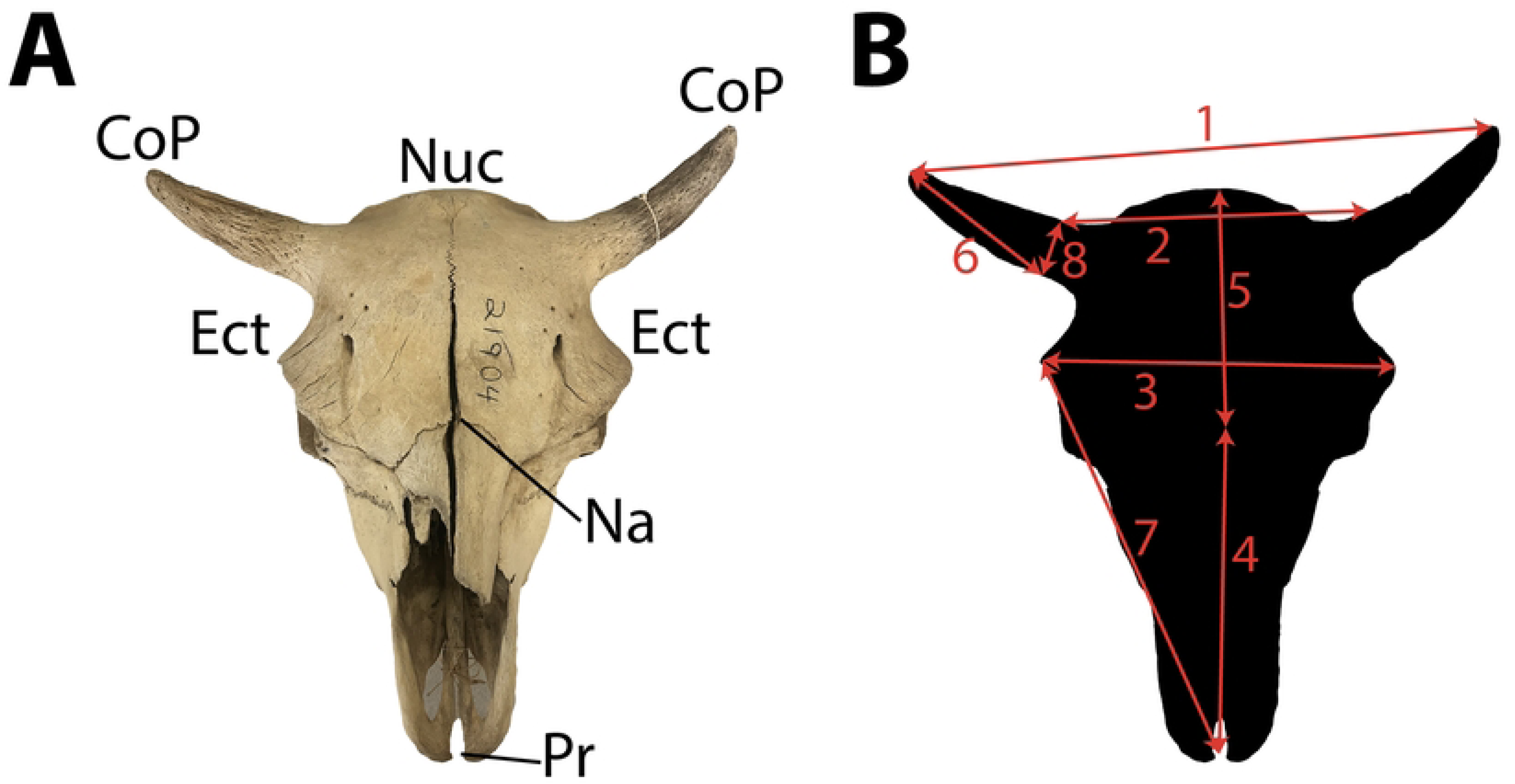
Cranial features and measurements referenced in study of *Bison bison*. (**A**) Relevant anatomical features on the skull of *Bison bison* (KU:Mamm 21904): CoP, Cornual process (frontal horn); Ect, Ectorbitale; Pr, Prosthion; Na, Nasion; Nus, Nuchal line. (**B**) Linear cranial measurements used in analyses: 1, Width of skull between tips of cornual processes; 2, Width of skull between bases of cornual processes; 3, Width of skull between left and right ectorbitale; 4, Length from prosthion to nasion; 5, Length from nasion to nuchal line; 6, Length of cornual process from base to tip; 7, Distance between prosthion and ectorbitale; 8, Width of cornual process at base. **Abbreviations**: KU:Mamm, mammalogy collections at the University of Kansas Natural History Museum.

## Results

### Univariate statistical tests

Tests of normality and unimodality demonstrate minimal evidence for sexual dimorphism in *U. anceps*. None of the eight cranial metric distributions deviated significantly (p ≤ 0.05) from normality, and only canine length showed a significantly non-unimodal distribution (Fig 5 and Table 1). In contrast, four metrics of interest (width between bases of frontal horns, width between left and right ectorbitale, length from nasion to nuchal line, and width of cornual process at base) in *B. bison* display distributions that differ significantly from normality (Fig 6 and Table 2). Other tests of normality and unimodality in *Bison* yield marginally nonsignificant results (0.05 < p ≤ 0.10).

**Figure 5.**
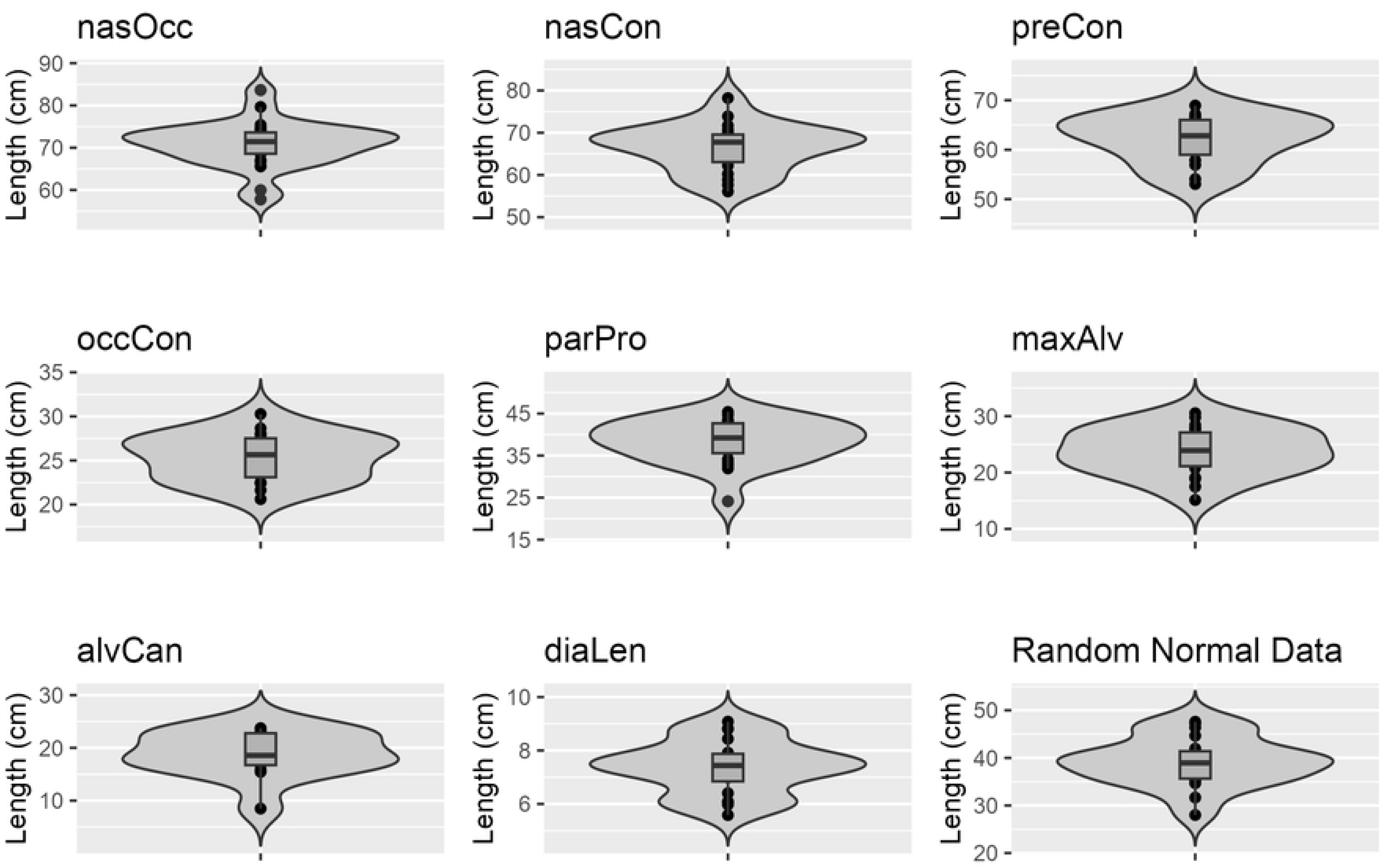
Distributions of cranial metrics in *Uintatherium anceps*. Random normal data generated with n, mean, and standard deviation equal to that of parPro. **Abbreviations:** nasOcc, length from nasals to occipital crest; nasCon, length from nasals to occipital condyles; preCon, length from premaxillaries to occipital condyles; occCon, height from occipital condyle to occipital crest; parPro, height of parietal protuberance; maxAlv, height of maxillary protuberance; alvCan, length of canine from alveolus to tip; diaLen, diastemal length.

**Table 1.** Results of Shapiro-Wilk and Hartigan’s dip tests in *Uintatherium anceps*.

|  | n | W Statistic | p-value | Dip Statistic | p-value |
| --- | --- | --- | --- | --- | --- |
| **Length from nasals to occipital crest** | 25 | 0.959 | 0.404 | 0.041 | 0.992 |
| **Length from nasals to occipital condyles** | 20 | 0.973 | 0.821 | 0.049 | 0.978 |
| **Length from premaxillaries to occipital condyles** | 15 | 0.935 | 0.329 | 0.067 | 0.841 |
| **Height from occipital crest to occipital condyle** | 20 | 0.968 | 0.703 | 0.073 | 0.507 |
| **Height from parietal protuberance to tip of postglenoid process** | 20 | 0.935 | 0.196 | 0.070 | 0.599 |
| **Height of maxillary protuberance** | 22 | 0.977 | 0.868 | 0.065 | 0.640 |
| **Length of canine from alveolus to tip** | 13 | 0.895 | 0.114 | 0.128 | 0.042* |
| **Diastemal length** | 20 | 0.960 | 0.542 | 0.072 | 0.535 |

W statistics (Shapiro-Wilk test for normality), dip statistics (Hartigan's dip test for unimodality), and associated p-values for the distributions of eight cranial metrics in *Uintatherium anceps*. Asterisks indicate significant p-values ( $\alpha = 0.05$ ).

**Figure 6.**
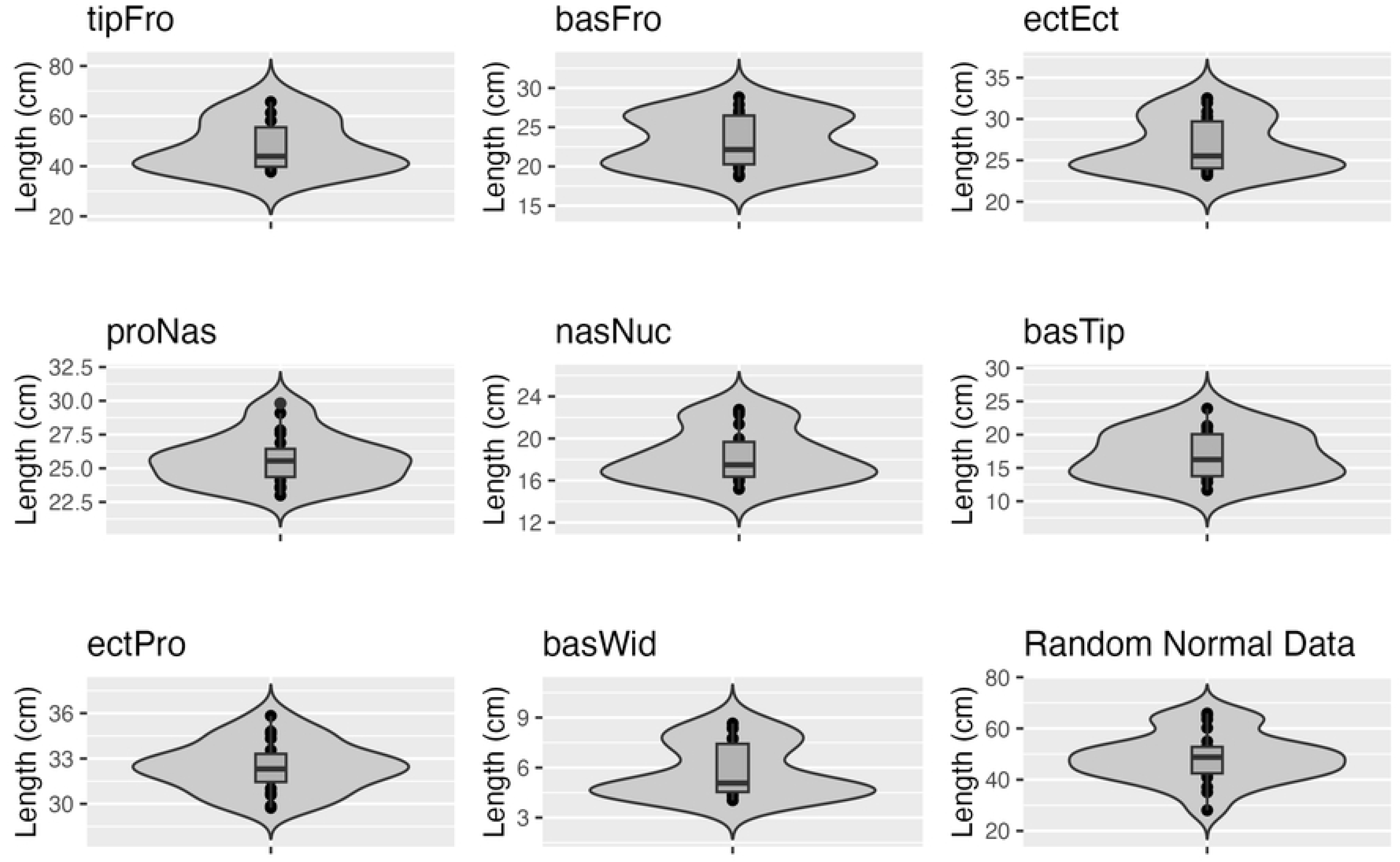
Distributions of cranial metrics in *Bison bison*. Random normal data generated with n, mean, and standard deviation equal to that of tipFro. **Abbreviations**: tipFro, width between tips of cornual processes (frontal horns); basFro, width between bases of cornual processes; ectEct, width between left and right ectorbitale; proNas, length from prosthion to nasion; nasNuc, length from nasion to nuchal line; basTip, length of cornual process; ectPro, length from prosthion to ectorbitale; basWid, width of cornual process at base.

**Table 2.** Results of Shapiro-Wilk and Hartigan’s dip tests in *Bison bison*.

|  | n | W Statistic | p-value | Dip Statistic | p-value |
| --- | --- | --- | --- | --- | --- |
| **Width between tips of cornual processes (frontal horns)** | 11 | 0.875 | 0.090 | 0.088 | 0.648 |
| **Width between bases of cornual processes** | 23 | 0.910 | 0.041* | 0.097 | 0.063 |
| **Width between left and right ectorbitale** | 23 | 0.871 | 0.007* | 0.062 | 0.703 |
| **Length from prosthion to nasion** | 20 | 0.954 | 0.431 | 0.063 | 0.782 |
| **Length from nasion to nuchal line** | 23 | 0.897 | 0.022* | 0.071 | 0.456 |
| **Length of cornual process** | 18 | 0.947 | 0.376 | 0.080 | 0.438 |
| **Length from prosthion to ectorbitale** | 20 | 0.979 | 0.914 | 0.066 | 0.702 |
| **Width of cornual process at base** | 23 | 0.840 | 0.002* | 0.078 | 0.296 |

W statistics (Shapiro-Wilk test for normality), dip statistics (Hartigan's dip test for unimodality), and associated p-values for the distributions of eight cranial metrics in *Bison bison*. Asterisks indicate significant p-values ( $\alpha = 0.05$ ).

### Mixture modeling

Mixture modeling analysis provides no evidence of sexual dimorphism in uintatheres. Among seven of the eight cranial metrics analyzed, the preferred model only had one component, with four components favored for one metric (Table 3). In contrast, among bison, two components were preferred for three of the selected cranial metrics (Table 4).

**Table 3.** Number of components preferred by mixture modeling for each cranial metric in Uintatherium anceps.

|  | Number of Components | BIC |
| --- | --- | --- |
| Length from nasals to occipital crest | 1 | -161.16444 |
| Length from nasals to occipital condyles | 1 | -130.53025 |
| Length from premaxillaries to occipital condyles | 1 | -94.51164 |
| Height from occipital crest to occipital condyle | 1 | -100.71799 |
| Height from parietal protuberance to tip of postglenoid process | 1 | -127.83192 |
| Height of maxillary protuberance | 1 | -129.91186 |
| Length of canine from alveolus to tip | 4 | -74.331029 |
| Diastemal length | 1 | -61.461477 |

Abbreviations: BIC, Bayesian Information Criterion.

**Table 4.** Number of components preferred by mixture modeling for each cranial metric in Bison bison.

|  | Number of Components | BIC |
| --- | --- | --- |
| Width between tips of cornual processes (frontal horns) | 4 | -82.300915 |
| Width between bases of cornual processes | 2 | -121.1501 |
| Width between left and right ectorbitale | 2 | -116.17363 |
| Length from prosthion to nasion | 1 | -86.664723 |
| Length from nasion to nuchal line | 3 | -110.16906 |
| Length of cornual process | 1 | -102.00116 |
| Length from prosthion to ectorbitale | 1 | -81.889981 |
| Width of cornual process at base | 2 | -76.239236 |

Abbreviations: BIC, Bayesian Information Criterion.

### Hierarchical clustering

Results of hierarchical clustering analysis do not favor previously hypothesized patterns of sexual dimorphism in uintatheres. After culling the dataset for imputation, 19 skulls remained, including three that have been considered females by previous authors and four that have been considered males [60,62,65]. Purported females and males occur apparently at random across the dendrogram and do not cluster at all according to their hypothesized sex (Fig 7). Likewise, putative females and males do not appear to differ consistently in morphology, as they vary widely in most metrics.

**Figure 7.**
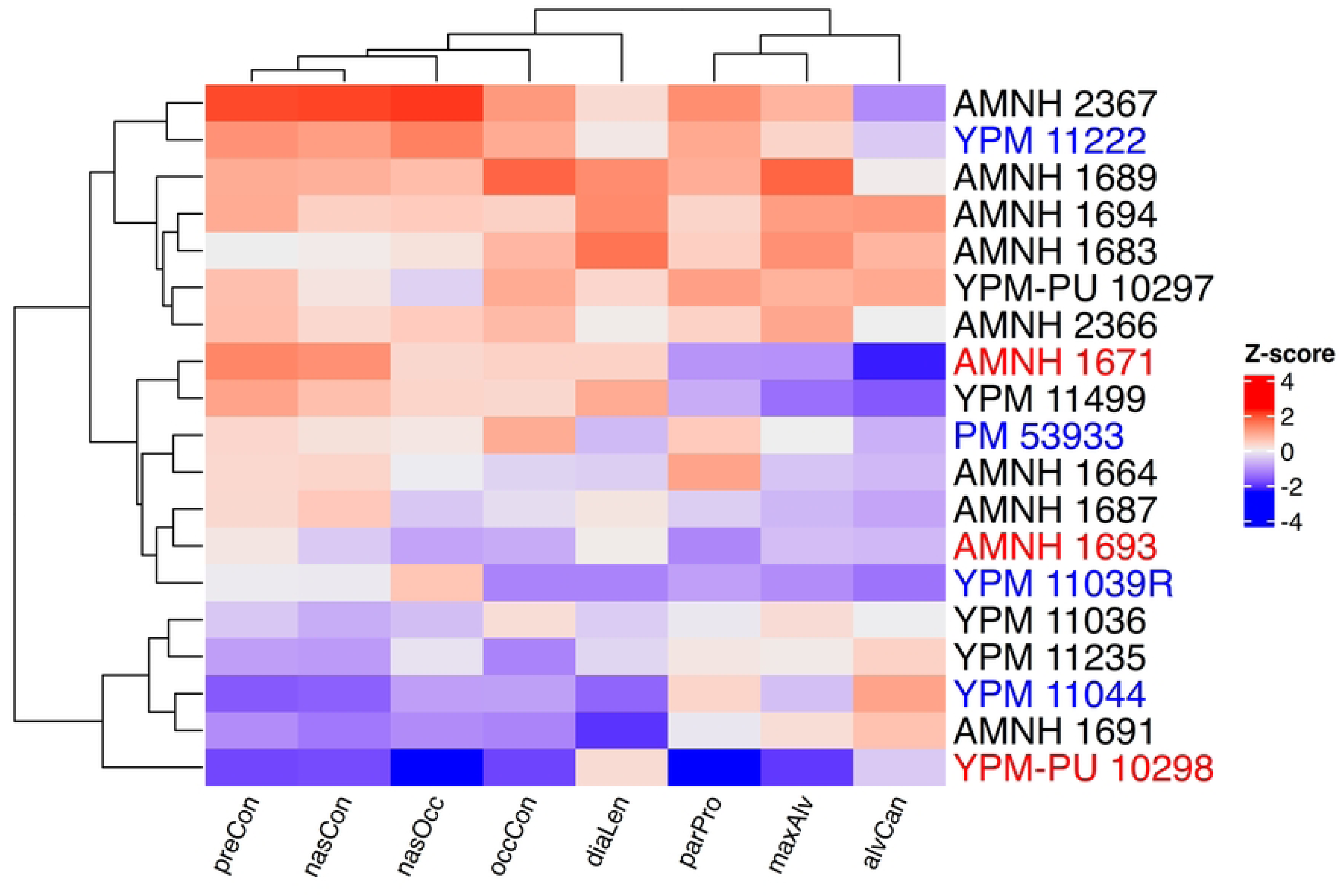
Hierarchical clustering of *Uintatherium anceps* crania. Skulls labeled in red and blue represent purported females and males according to previous authors, respectively. **Abbreviations**: nasOcc, length from nasals to occipital crest; nasCon, length from nasals to occipital condyles; preCon, length from premaxillaries to occipital condyles; occCon, height from occipital condyle to occipital crest; parPro, height of parietal protuberance; maxAlv, height of maxillary protuberance; alvCan, length of canine from alveolus to tip; diaLen, diastemal length.

In contrast, results of hierarchical clustering in *Bison* are consistent with sexual dimorphism. Bison skulls form two primary clusters, and with only one exception, all females fall in one cluster while all males fall in the other (Fig 8). Males are consistently larger than females, especially in metrics related to the size of the cornual processes and the width of the skull.

**Figure 8.**
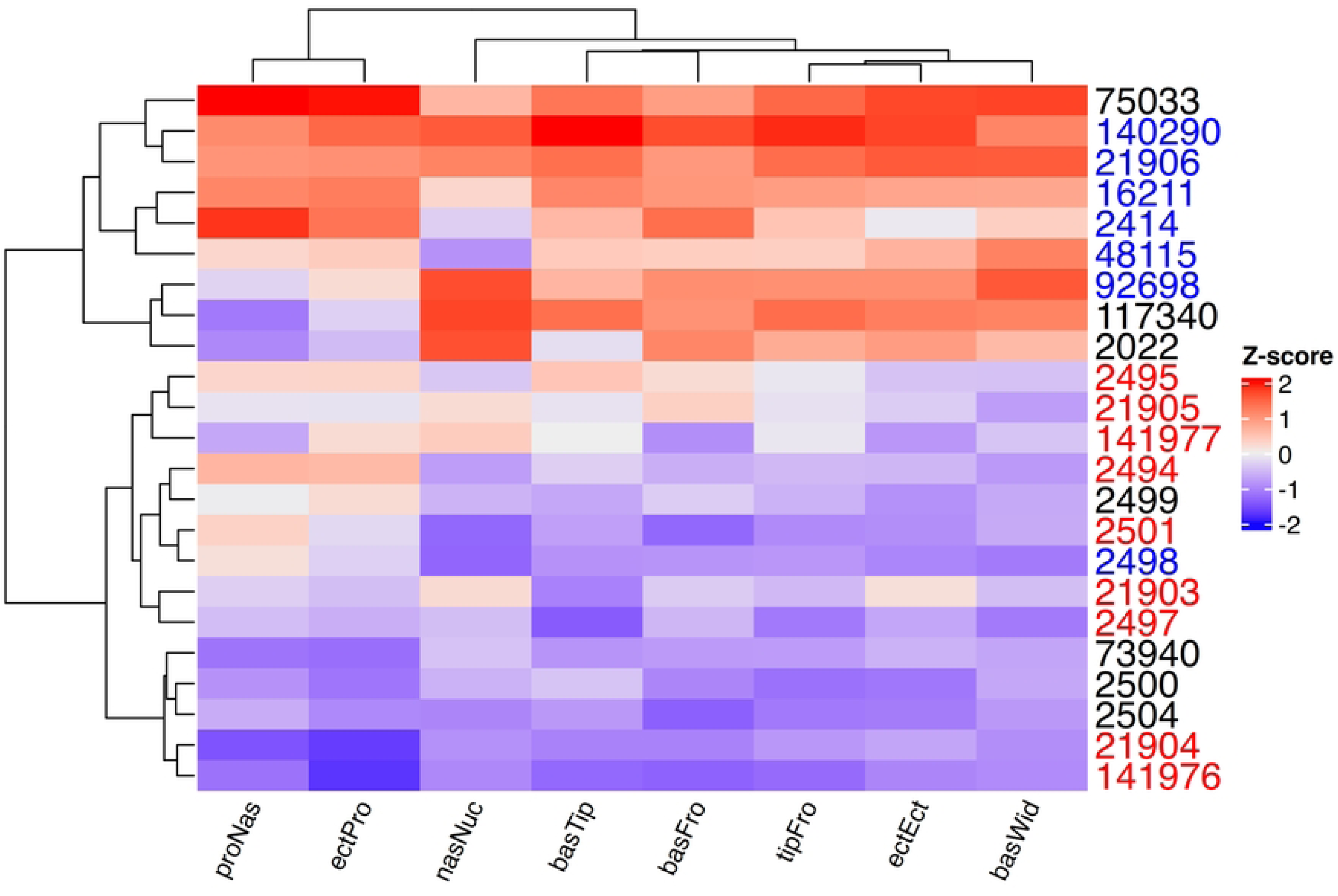
Hierarchical clustering of *Bison bison* skulls. Skulls labeled in red and blue represent confirmed females and males, respectively. **Abbreviations**: tipFro, width between tips of cornual processes (frontal horns); basFro, width between bases of cornual processes; ectEct, width between left and right ectorbitale; proNas, length from prosthion to nasion; nasNuc, length from nasion to nuchal line; basTip, length of cornual process; ectPro, length from prosthion to ectorbitale; basWid, width of cornual process at base. All specimens curated at the University of Kansas Natural History Museum (KU:Mamm).

### Principal component analysis and k-means clustering

Results of PCA are not immediately indicative of sexual dimorphism in *U. anceps*. Together, PC1 and PC2 account for over 80% of observed variance, about 59% and 25%, respectively (S6 Table). PC1 is positively correlated with all variables except for canine length, while PC2 is highly positively correlated with the length of the canine and the height of the maxillary and parietal horns, but negatively correlated with variables capturing the total length of the skull (S7 Table). Thus, PC1 seems to reflect the overall size of the skull while PC2 mostly captures the relative prominence of the horns and canine. PC3 is only highly positively correlated with diastemal length. All purported female specimens show low to moderate PC1 and PC2 values; purported males vary widely in their PC1 and PC2 values (Fig 9A)..

**Figure 9.**
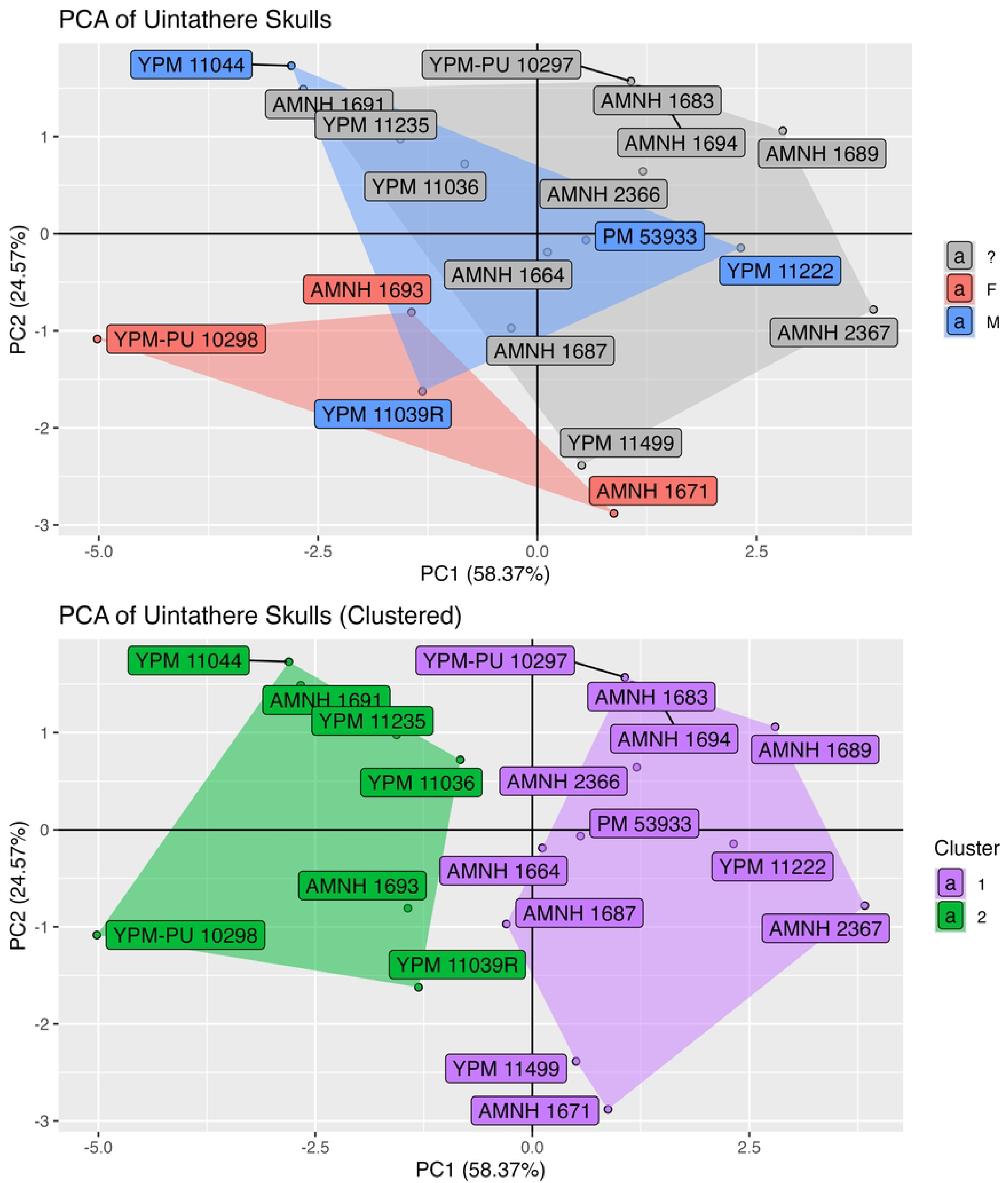
Results of PCA in *Uintatherium anceps*. (**A**) Bivariate plot displaying values of PC1 and PC2 for each specimen of *Uintatherium anceps*. Specimens labeled in red represent purported females, blue specimens are purported males, and gray specimens have not been attributed to either sex by previous authors. (**B**) Same bivariate plot but with specimens labeled in green and purple belonging to cluster 1 and 2, respectively, as designated by k-means clustering.

Taken together with results of k-means cluster analysis, PCA does not provide evidence of sexual dimorphism in *Uintatherium*. The two most favorable clusters (n=12 and n=7) are separated primarily along the PC1 axis, and both include a mix of purported females and males (Fig 9B). Of the three methods employed to determine the optimal number of clusters, none favor two (S2 Fig). The gap statistic favors one cluster, while the total within sum of squares and silhouette methods indicate six. Fisher’s Exact Test indicates that purported sex and k-means cluster assignment are not correlated (p=1.0).

Results of PCA and k-means cluster analysis are consistent with sexual dimorphism in *Bison*. PC1 and PC2 are the only significant (>5% of total variance) components, composing about 78% and 15% of the variance, respectively (S8 Table). All variables contribute positively to PC1, while only the variables capturing the length of the anterior portion of the skull contribute positively to PC2 (S9 Table). All females have negative PC1 scores, while all but one male have positive PC1 scores (Fig 10A). Both sexes vary in their PC2 scores. When the two most favorable clusters are formed through k-means clustering, all females are assigned to the same cluster, while all males except for one fall into the other (Fig 10B). Fisher’s exact test indicates significant correlation between sex and assigned cluster (p = 0.029). All three methods implemented to determine the optimal number of clusters favor two (S3 Fig).

**Figure 10.**
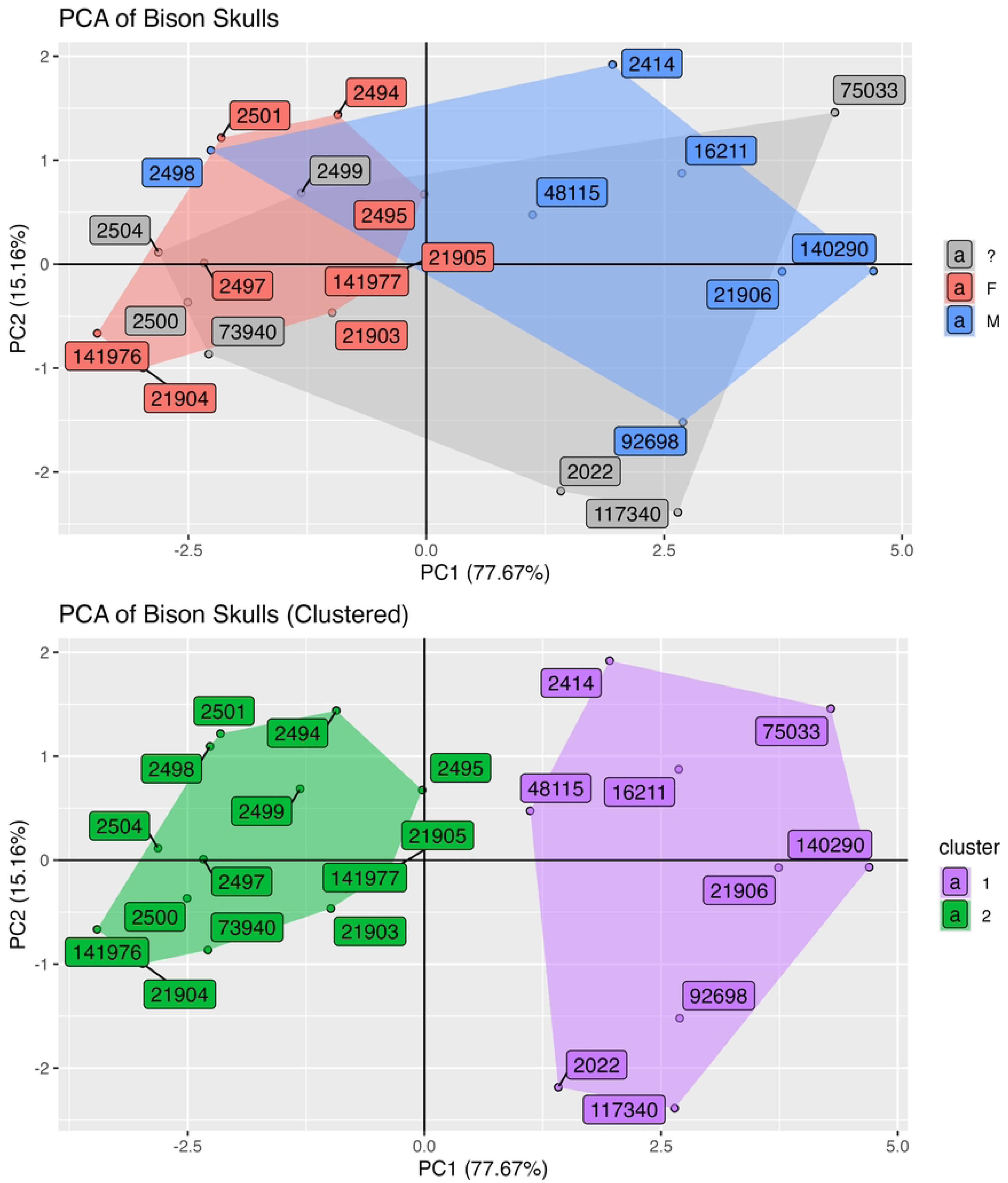
Results of PCA in *Bison bison*. (**A**) Bivariate plot displaying values of PC1 and PC2 for each specimen of *Bison bison*. Specimens labeled in red represent confirmed females, blue specimens are confirmed males, and gray specimens lack sex metadata. (**B**) Same bivariate plot but with specimens labeled in green and purple belong to cluster 1 and 2, respectively, as designated by k-means clustering. All specimens curated at the University of Kansas Natural History Museum (KU:Mamm).

### Effect size statistics

Effect size statistics may provide some evidence of sexual dimorphism in *Uintatherium*. Maxillary horn height and canine length exhibit larger effect sizes than metrics capturing the overall length of the skull (Fig 11 and S10 Table). These results are consistent with the expectation that features directly under sexual selection would exhibit higher effect sizes than other traits. That said, the residual-based sex assignments of individual specimens do not align with the hypotheses of previous authors as to which individuals are male and female. Furthermore, confidence intervals (CI) and prediction intervals (PI) overlap in most metrics.

**Figure 11.**
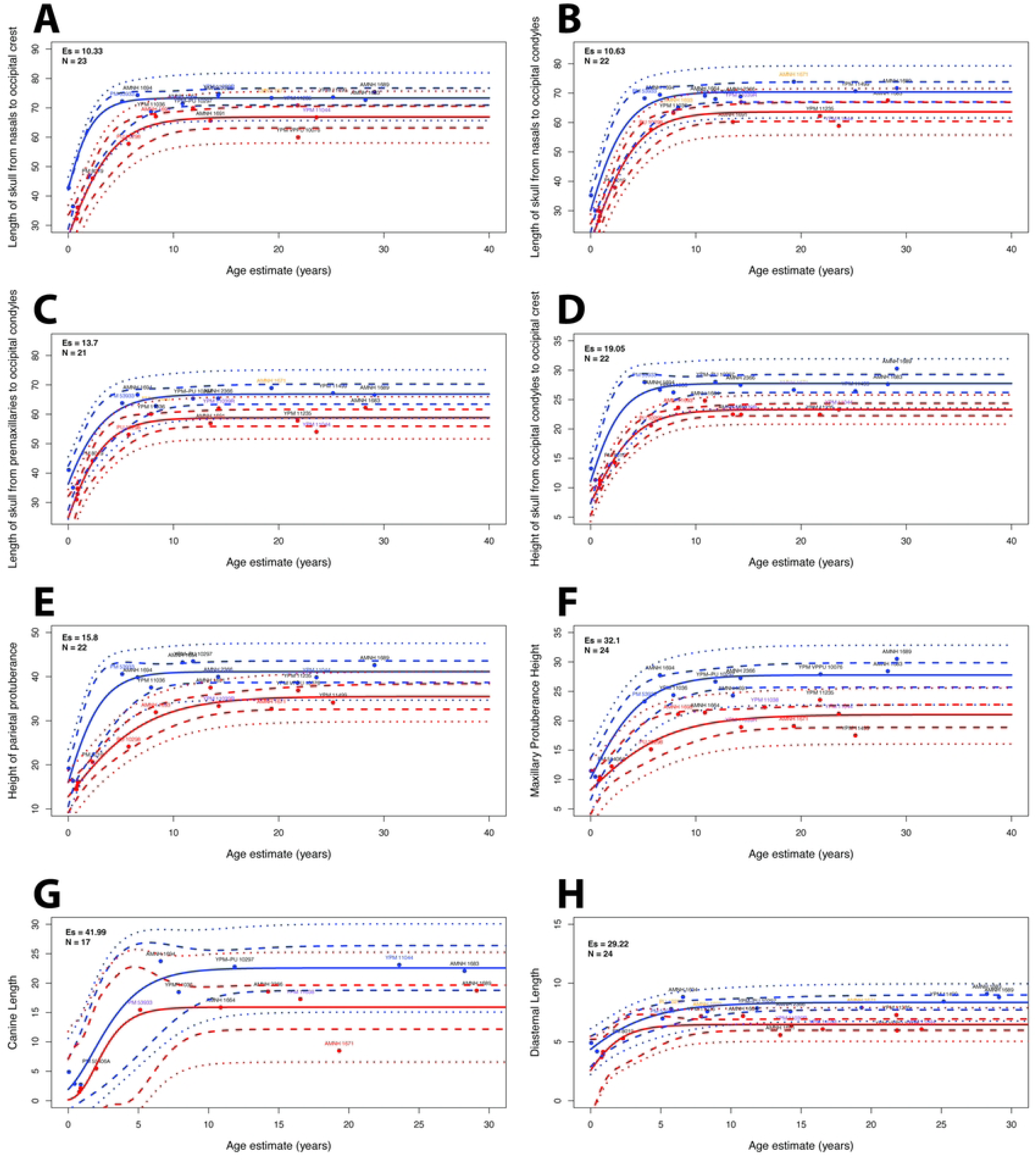
Results of effect size analysis in *Uintatherium anceps*. Sex-specific Gompertz regression curves for eight cranial metrics in *Uintatherium anceps*: (**A**), length of skull from nasals to occipital crest; (**B**), length of skull from nasals to occipital condyles; (**C**), length of skull from premaxillaries to occipital condyles; (**D**), height of skull from top of occipital crest to bottom of occipital condyle; (**E**), height of skull from top parietal protuberance to bottom of postglenoid process; (**F**), height of maxillary protuberance; (**G**), canine length; (**H**), diastemal length. Sexes assigned using residual method. 95% Confidence intervals (dashed lines) and prediction intervals (dotted lines) shown for each sex. Specimens labeled in red represent purported females with negative residual scores, specimens labeled in orange are purported females with positive residual scores, specimens labeled in blue are purported males with positive residual scores, and specimens labeled in purple are purported males with negative residual scores. **Abbreviations**: Es, effect size; N, number of specimens (including simulated “newborn” data).

Effect size statistics indicate a more extreme degree of sexual dimorphism in *Bison*. Traits related to horn size exhibit higher effect sizes than those correlated with the overall size of the skull, highlighting the effects of sexual selection (Fig 12 and S11 Table). The highest effect sizes are greater than those observed in any metric in *Uintatherium*, as is the disparity between more and less dimorphic traits. While there are instances where residual-based sex assignment is incorrect in *Bison*, PIs and CIs are generally better separated than in *Uintatherium*.

**Figure 12.**
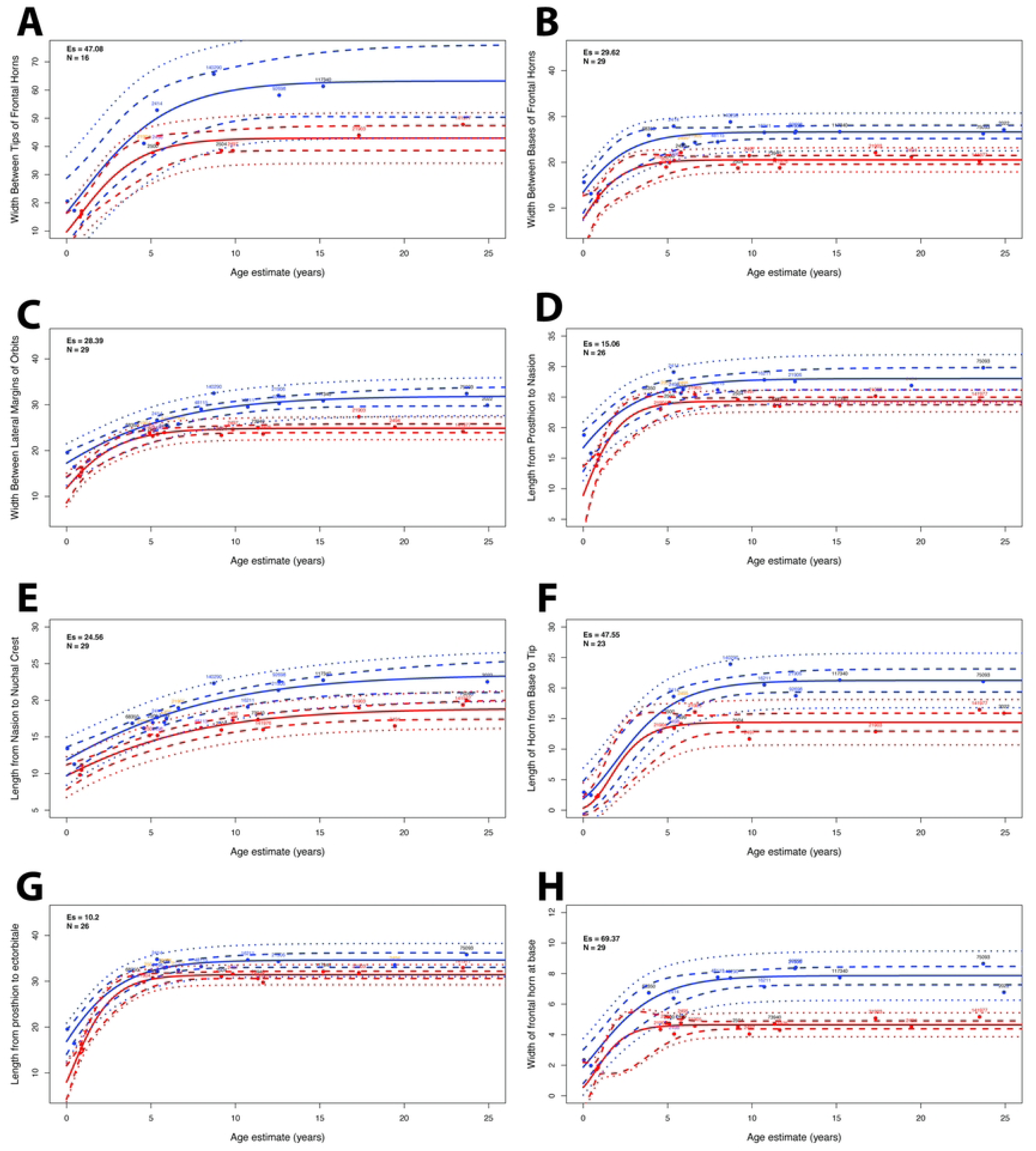
Results of effect size analysis in *Bison bison*. Sex-specific Gompertz regression curves for eight cranial metrics in *Bison bison*: (**A**), width of skull between tips of cornual processes; (**B**), width of skull between bases of cornual processes; (**C**), width of skull between left ectorbitale and right ectorbitale; (**D**), length from prosthion to nasion; (**E**), length from nasion to nuchal line; (**F**), length of cornual process from base to tip; (**G**), length from ectorbitale to prosthion; (**H**), width of cornual process at base. Sexes assigned using residual method. 95% Confidence intervals (dashed lines) and prediction intervals (dotted lines) shown for each sex. Specimens labeled in red represent females with negative residual scores, specimens labeled in orange are females with positive residual scores, specimens labeled in blue are males with positive residual scores, and specimens labeled in purple are males with negative residual scores. **Abbreviations**: Es, effect size; N, number of specimens (including simulated “newborn” data).

## Discussion

Of all approaches employed, none provide evidence for *an extreme degree of sexual dimorphism* in *Uintatherium* [65] (emphasis added by author). Results are generally indistinguishable from monomorphy or suggest only moderate sexual dimorphism. In contrast, the same approaches reveal clear signals of strong, consistent sexual dimorphism in *Bison*. These findings indicate that *U. anceps* may have been less sexually dimorphic than previous authors have postulated, or at least that evidence of strong sexual dimorphism is currently lacking in this taxon.

Statistical tests fail to reject normality and unimodality in all uintathere metrics except canine length (Table 1). While there are more significant results in *Bison* (Table 2), failure to reject unimodality and normality in several known dimorphic traits reaffirms previous authors findings that classical univariate statistical tests lack sufficient power to detect sexual dimorphism in most cases. Univariate mixture modeling appeared to perform marginally better at detecting dimorphism than tests of normality and unimodality, but these results were also ambiguous (Tables 3 and 4).

More informative were the results of multivariate analyses. In hierarchical clustering of *Bison*, skulls were split into two primary clusters, which were highly correlated with sex (p=0.001) (Fig 8). However, in *Uintatherium*, purported females and males were distributed widely and unpredictably across the dendrogram (Fig 7). This suggests either that hierarchical clustering in the current sample of *Uintatherium* crania does not correlate with sex or that previous authors have been incorrect in their interpretation of which specimens belong to which sex. This finding is corroborated by results of PCA and k-means cluster analysis, which suggest that purported male and female uintatheres cannot be distinguished by traditional morphometric approaches (Fig 9). The favoring of two clusters in *Bison* (S3 Fig), and the fact that these two clusters are highly correlated with sex (Fig 10), further suggest that the statistical signal of dimorphism in *Uintatherium* is weaker than it is in *Bison*.

Effect size statistics, especially considered in light of analyses in *Bison*, also suggest a relatively weak signal of dimorphism in *Uintatherium*. Purportedly dimorphic traits, especially the size of the canine and maxillary horns, did show higher effect sizes than characters expected to be relatively monomorphic (Fig 11 and S10 Table). These results indicate that *Uintatherium* may have shown some degree of dimorphism in the expression of these characters. However, the residual-based sex assignment used to generate these effect sizes did not align with previous authors’ interpretations of the sex of individual specimens (Fig 11)[60,62,65]. The discrepancy in effect size between more and less dimorphic traits in *Bison* was more extreme than in *Uintatherium*, as was the effect size of the most dimorphic characters (Figs 11 and 12, S10 and S11 Tables). As such, even if *Uintatherium* did exhibit some degree of sexual variation, the current evidence suggests that it was less dimorphic than *Bison*.

It is possible that the linear metrics chosen and statistical analyses performed failed to provide sufficient power to detect sexual dimorphism. CT scanning uintathere skulls may allow for gathering of more precise measurements and application of three-dimensional geometric morphometric approaches that could reveal morphological patterns undetected herein. However, the unambiguous signal of sexual dimorphism detected in a similar suite of linear metrics gathered from a comparable sample of *Bison* suggests that traditional, linear morphometrics can provide the necessary power to detect sexual dimorphism without a priori knowledge of individual sex.

Heavily biased sex ratios have been documented elsewhere in the fossil record, and it is possible that a bias towards male specimens, as has been posited by previous authors, may obscure the statistical signal of sexual dimorphism in *Uintatherium* [48,62,65,97,98]. However, research has shown that male and female mammals are subject to approximately equivalent postmortem preservation, making a taphonomic bias towards male uintatheres unlikely [98]. Collection bias towards larger, more robust specimens with prominent horns has been proposed as another potential explanation for disproportionate representation of males among the current sample of uintatheres, but, given both sexes’ immense size and bizarre appearance in comparison to coeval mammals, I find this argument uncompelling [62,63,97]. Furthermore, while male-biased sex ratios are well-documented in the mammalian fossil record, confirmed instances have been largely confined to polygynous species of herding mammals [97–99]. Such taxa are often characterized by wide-ranging males that are more broadly distributed across a landscape, and therefore more likely to fossilize [97–99]. Unlike other similarly sized and contemporaneous mammals, however, no uintathere death assemblage has ever been recovered, and thus there is no compelling evidence that they lived in polygynous herds [88,96,100]. Given the warm, wet, rugged, and heavily forested landscape in which uintatheres appear to have lived, it is perhaps more reasonable to infer that uintatheres were largely solitary, and that females and males were subject to similar spatial distributions [67,101–104]. I therefore see no reason to assume that the current sample of *Uintatherium* skulls is heavily skewed towards males.

I do not argue that these results allow us to conclusively reject the hypothesis that *U. anceps* was sexually dimorphic, nor do I claim that I have unequivocally shown that the taxon was totally or nearly monomorphic. Instead, I believe that these results allow us to refute the classical interpretation that the species exhibits evidence of “extreme” sexual dimorphism [60,62,65]. My results show that previous conceptualizations of sexual dimorphism in *Uintatherium* lack statistical support. Moreover, I find less evidence for sexual variation in *Uintatherium* than in *Bison*. Given their unresolved phylogenetic affinities and the clade-specific nature of sexual dimorphism among extant mammals, there is little justification for assuming that uintatheres were strongly sexually dimorphic a priori. Furthermore, large herbivorous extant mammals with apparently similar ecological preferences have been shown to exhibit limited degrees of dimorphism [7,105]. This analysis provides the first quantitative data in the discussion of dimorphism among uintatheres, and these results suggest that these animals were not strongly dimorphic.

These results do not preclude the presence of sexual dimorphism in Dinocerata altogether; to the contrary, they provide the first quantitative evidence of modest dimorphism in uintathere canine length and horn size (Figs 5 and 11, Table1, S10 Table). Traits not examined in this study, such as the inframandibular flange, may have been subject to different patterns of sexual variation. Patterns of sexual dimorphism may have also varied between uintathere taxa, as in extant mammalian orders [106]. Nonetheless, these results suggest that reinterpretation of uintatheres, particularly *U. anceps*, as only slightly or moderately dimorphic represents a viable hypothesis in light of currently available data. Only by analyzing additional taxa, applying more robust morphometric techniques, and uncovering additional fossil specimens can we make further progress in assessing sexual dimorphism among uintatheres. Given the potentially relatively basal position of uintatheres on the Placental phylogeny, these results may also lend credence to the hypothesis that strong sexual dimorphism was not ancestral among mammals.

The linear morphometric approach employed here is easily reproducible and broadly applicable in the study of sexual dimorphism among extinct animals. I therefore advocate for the application of these techniques in analyses of sexual dimorphism among other fossil clades.

Through a quantitative, comparative framework, we may begin to understand when and how many times dimorphism has evolved across the tree of life.

## Acknowledgements

I would like to thank K. Christopher Beard for reviewing this article and for his help in formulating this project. I thank Bruce Lieberman for his input on analyses and data visualization. I also thank Kelly Matsunaga, Mark Holder, Marina Suarez, and Jamie Walters for various discussions and suggestions. I thank Geoff Flora for correspondence and insights about uintathere biology. Lastly, I am grateful to Judy Galkin at the American Museum of Natural History, Vanessa Rhue and Dan Brinkman at the Yale Peabody Museum, and Dianna Krejsa and Jocelyn Colella at the University of Kansas Biodiversity Institute and Natural History Museum for access to specimens.

## Supporting information

**S1 Table. Uintathere cranial measurements, abbreviations, and definitions as used in this study.** All measurements obtained in ImageJ.

**S2 Table. Tests for measurement error in *Uintatherium anceps* cranial metrics.** Results of tests for association using Pearson’s product moment correlation coefficient evaluating the relationship between the mean of hand-measured and ImageJ values and the difference between hand-measured and ImageJ values for each cranial metric in *Uintatherium anceps*. n indicates the number of specimens for which values were obtained both by hand and by ImageJ for each metric. No tests were significant (⍺ = 0.05).

**S3 Table. Age grades, their numeric ranges, and definitions implemented in effect size statistical analyses of *Uintatherium anceps*.**

**S4 Table. Bison cranial measurements, abbreviations, and definitions as used in this study.** All measurements obtained in ImageJ.

**S5 Table. Age grades, their numeric ranges, and definitions implemented in effect size statistical analyses of *Bison bison***. Adapted from Fuller [85].

**S6 Table. Eigenvalues of and variance explained by each component in principal component analysis of *Uintatherium anceps* cranial metrics**. Abbreviations: PC, principal component.

**S7 Table. Principal component loadings of cranial metrics in *Uintatherium anceps***. Abbreviations: PC, principal component.

**S8 Table. Eigenvalues of and variance explained by each component in principal component analysis of *Bison bison* cranial metrics**. Abbreviations: PC, principal component.

**S9 Table. Principal component loadings of cranial metrics in *Bison bison*.** Abbreviations: PC, principal component.

**S10 Table. Effect sizes in *Uintatherium anceps* cranial metrics.** Number of specimens and effect sizes for each metric of interest in *Uintatherium anceps*. **Abbreviations**: n, number of specimens after simulation of young individuals; n*, number of specimens before simulation of young individuals; ES, effect size.

**S11 Table. Effect sizes in *Bison bison* cranial metrics.** Number of specimens and effect sizes for each metric of interest in *Bison bison*. **Abbreviations**: n, number of specimens after simulation of young individuals; n*, number of specimens before simulation of young individuals; ES, effect size.

**S1 Figure. Bland-Altman plots for each cranial metric in *Uintatherium anceps*.** Plots display the relationship between the mean of hand-measured and ImageJ values and the difference between hand-measured and ImageJ values for each cranial metric in *Uintatherium anceps*. (A), length of skull from nasals to occipital crest; (B), length of skull from nasals to occipital condyles; (C), length of skull from premaxillaries to occipital condyles; (D), height of skull from top of occipital crest to bottom of occipital condyle; (E), height of skull from top parietal protuberance to bottom of postglenoid process; (F), height of maxillary protuberance; (G), canine length; (H), diastemal length.

**S2 Figure. Number of clusters present in *Uintatherium anceps* crania.** Optimal number of clusters in *Uintatherium* data as indicated by total within sum of squares, average silhouette width, and gap statistic.

**S3 Figure. Number of clusters present in *Bison bison* crania.** Optimal number of clusters in *Bison* data as indicated by total within sum of squares, average silhouette width, and gap statistic.

**S1 File. *Uintatherium anceps* cranial data used in traditional morphometric analyses (partial).** This file only includes American Museum and Yale Peabody Museum specimens.

**S2 File. *Uintatherium anceps* cranial data used in traditional morphometric analyses (complete).** This file includes American Museum and Yale Peabody Museum specimens as well as data from Turnbull [65] for Field Museum skulls.

**S3 File. *Bison bison* cranial data used in traditional morphometric analyses.**

**S4 File. *Uintatherium anceps* cranial data used in effect size analyses.**

**S5 File. *Bison bison* cranial data used in effect size analyses.**

**S6 File. Measurements of uintathere skulls gathered by hand.** Used in tests for correlation between skull size and measurement error.

**S7 File. Measurements of uintathere skulls obtained through ImageJ.** Used in tests for correlation between skull size and measurement error.

**S8 File. Code used in traditional morphometric analyses of uintathere crania.**

**S9 File. Code used in traditional morphometric analyses of bison crania.**

**S10 File. Code used in effect size analyses of uintathere crania.**

**S11 File. Code used in effect size analyses of bison crania.**

**S12 File. Code used in tests of correlation between uintathere skull size and measurement error.**

